# Identification of AAV Capsids with Enhanced Intravitreal Transduction and Favorable Safety Profile in Non-Human Primates

**DOI:** 10.64898/2026.04.23.720493

**Authors:** Zhenhua Wang, Hao Li, Zhen Sun, Xianggang Xu, Qun Zhang, Peilu Zhao, Long Wang, Tongqian Xiao, Mengmeng Yu, Shengwen Wang, Rui He, Can Hu, Daoyuan Li, Bo Sun, Linlin Zhang, Youguang Luo, Zhenming An

**Author notes:** These authors contributed equally (Zhenhua Wang, Hao Li, Zhen Sun, Xianggang Xu). Correspondence: Youguang Luo,; Zhenming An,.

## Abstract

We report the discovery of novel adeno-associated virus (AAV) capsid variants engineered for superior intravitreal (IVT) gene delivery to the primate retina. Utilizing the REACH platform, we constructed a diverse AAV variant library and employed a multi-stage screening strategy involving in vitro selection on human retinal pigment cells followed by direct in vivo screening in non-human primates (NHPs). Following IVT administration in NHPS of a barcoded variant pool, next-generation sequencing analysis of retinal tissues identified lead candidates (e.g., E52, E54, and E57) that achieved transduction levels in the neural retina and RPE 5-10 fold higher than the benchmark R100. Concurrently, these high-potency variants exhibited an exceptional ocular confinement profile, with minimal to undetectable vector genome distribution in systemic organs. This combination of markedly enhanced retinal transduction and stringent local tropism establishes these engineered capsids as promising next-generation vectors for the treatment of inherited and acquired retinal diseases via a minimally invasive IVT route.

## INTRODUCTION

Inherited retinal diseases are a leading cause of irreversible vision loss and blindness.^1^ Furthermore, prevalent retinal disorders such as neovascular age-related macular degeneration (nAMD) and diabetic macular edema (DME) impose a significant public health burden due to their high prevalence and requirement for long-term, repeated treatments. Gene therapy, particularly strategies based on adeno-associated virus (AAV) vectors for gene replacement, gene editing, or sustained protein delivery, offers revolutionary therapeutic potential for these conditions.^2,3^ However, the efficiency of intraocular targeted delivery is paramount to the success and safety of such therapies.^3,4^ Among various administration routes, intravitreal (IVT) injection presents considerable clinical translational potential due to its relative simplicity, lower invasiveness compared to subretinal injection, and ability to broadly target the vitreous and inner retinal layers. The core challenge remains enabling therapeutic genes to efficiently and safely penetrate intraocular barriers (especially the inner limiting membrane, ILM) following IVT administration and to be delivered precisely to specific target cells—such as photoreceptors, retinal pigment epithelium (RPE), and ganglion cells—while minimizing off-target effects.^3–6^

Current engineering of AAV capsids for the IVT route has made progress. Traditional serotypes (e.g., AAV2, AAV8) often exhibit low transduction efficiency post-IVT due to the ILM barrier and interactions with vitreal proteins, particularly failing to reach the outer retinal cells, such as RPE, frequently requiring high vector doses for efficacy. High doses increase risks of intraocular inflammation and photoreceptor stress and may lead to systemic vector leakage.^7,8^ Directed evolution has yielded novel capsids like AAV2.7m8 and R100, showing enhanced photoreceptor transduction in mice and NHPs.^9,10^ However, translating these into human clinics demands further improvements in comprehensive performance: enhancing transduction efficiency to lower doses and reduce local inflammation; reducing off-target tropism to minimize systemic leakage; and improving manufacturability metrics like yield and stability.

In this study, we employed our proprietary REACH platform, constructing a highly diverse AAV capsid library. Through multiple rounds of in vitro and in vivo screening, engineered AAV variant capsids demonstrated significantly enhanced transduction efficiency in target tissues (neural retina or RPE), high safety profile, and promising developability, indicating substantial clinical translation potential.

## RESULTS

### 1. Proprietary REACH AAV Delivery Platform

Wild-type AAVs (such as AAV2, AAV8) are preferred vectors for ocular gene therapy due to their favorable safety profile and ability to transduce retinal cells. However, a key limitation is their poor efficiency in infecting outer retinal photoreceptors and RPE cells via the clinically simpler IVT route, often necessitating more invasive subretinal injection. To overcome this, we utilized our in-house proprietary REACH platform to generate a high-diversity AAV variant library. A multi-tiered screening system (in vitro screening, in vivo screening, and lead confirmation) was implemented, ultimately yielding engineered candidate vectors with high retinal penetration potency (Figure 1).

**Figure 1.**
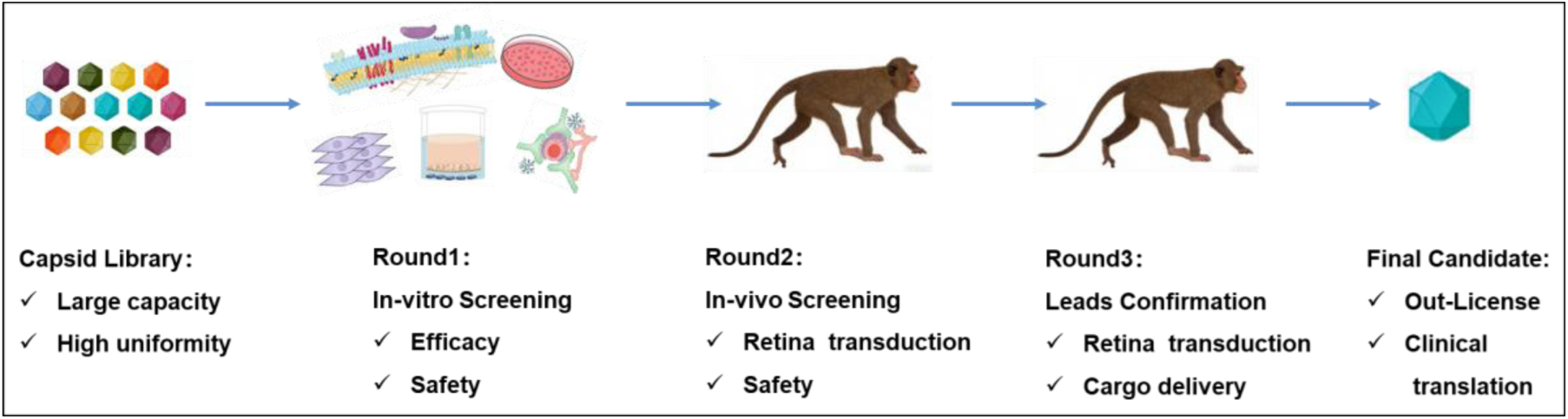
Schematic of the engineered AAV capsid screening pipeline based on REACH platform. A high-capacity, highly uniform AAV library was constructed de novo. The primary screening was performed through in vitro validation, yielding a lead library for next screening. This enriched library was then administered intravitreal to non-human primates (NHPs) for a secondary in vivo screening round, leading to the identification of a series of highly efficient retina-transduction AAV variants. The top-performing variants were further characterized individually with a therapeutic cargo in NHPs to get more comprehensive efficacy and safety profile, establishing a foundation for subsequent clinical translation.

### 2. Engineered Variants Demonstrate Significantly Enhanced Transduction of Human RPE Cells in Vitro

As RPE cells are crucial for visual function and a key target for gene therapy, we used ARPE-19 cells to evaluate the RPE transduction efficiency of AAV variants. Results showed that selected lead variants (E15, E35, E43, E52, E54, E57, E65) exhibited significantly enhanced infection efficiency and fluorescence intensity in ARPE-19 cells compared to the wild-type AAV2 control, also outperforming the AAV2.7m8 benchmark (Figure 2A-2D). Evaluation in HEK293 cells as a non-target cell model showed similar and comparable infection efficiency among variants, AAV2, and AAV2.7m8, suggesting the engineered variants possess certain infection specificity (Figure 2E-2H).

**Figure 2.**
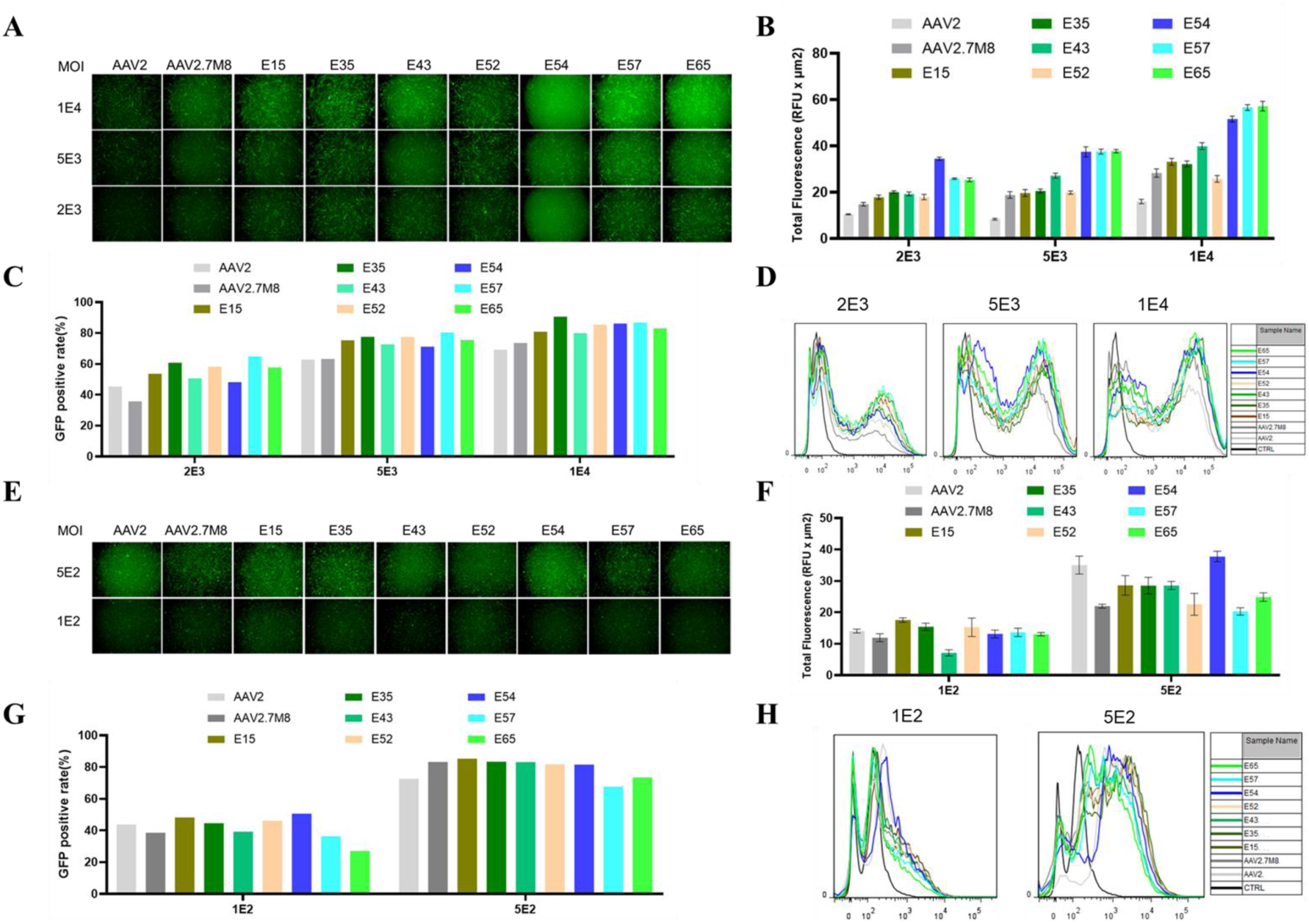
Engineered variants show significantly enhanced transduction of ARPE19 cells. (A) Representative fluorescence images of ARPE19 cells. AAV2 and corresponding variants expressing EGFP were used to infect ARPE19 cells in different MOIs (2E3、5E3 and 1E4) for 48 hours, and then fluorescence images were taken by microscopy. (B) Fluorescence intensity analysis of the figures as described in A. (C) Flow cytometry analysis of the percentage of GFP-positive ARPE19 cells. AAV2 and corresponding variants expressing EGFP were used to infect ARPE19 cells in different MOIs (2E3、5E3 and 1E4) for 48 hours, and then these cell samples were collected and analyzed by flow cytometry. (D) Flow cytometry analysis of fluorescence intensity in ARPE19 cells. Experiments were performed as described in C, and then the fluorescence intensity of each sample was analyzed. (E) Representative fluorescence images of HEK293 cells. AAV2 and corresponding variants expressing EGFP were used to infect HEK293 cells in different MOIs (1E2 and 5E2) for 48 hours, and then fluorescence images were taken by microscopy. (F) Fluorescence intensity analysis of the figures as described in E. (G) Flow cytometry analysis of the percentage of GFP-positive HEK293 cells. AAV2 and corresponding variants expressing EGFP were used to infect HEK293 cells in different MOIs (1E2 and 5E2) for 48 hours, and then these cell samples were collected and analyzed by flow cytometry. (H) Flow cytometry analysis of fluorescence intensity in HEK293 cells. Experiments were performed as described in G, and then the fluorescence intensity of each sample was analyzed.

### 3. Engineered Variants Shows Superior Retina transduction in Non-Human Primates

Samples for NHP studies were produced at a 2-3L bioreactor scale, purified via TFF, affinity chromatography, and density gradient centrifugation, achieving an overall yield of 29.2%. Quality assessments confirmed high purity, >70% full capsid ratio, homogeneity (DLS average size: 24.99 nm), and acceptable endotoxin levels (Figure 3A-3C).

**Figure 3.**
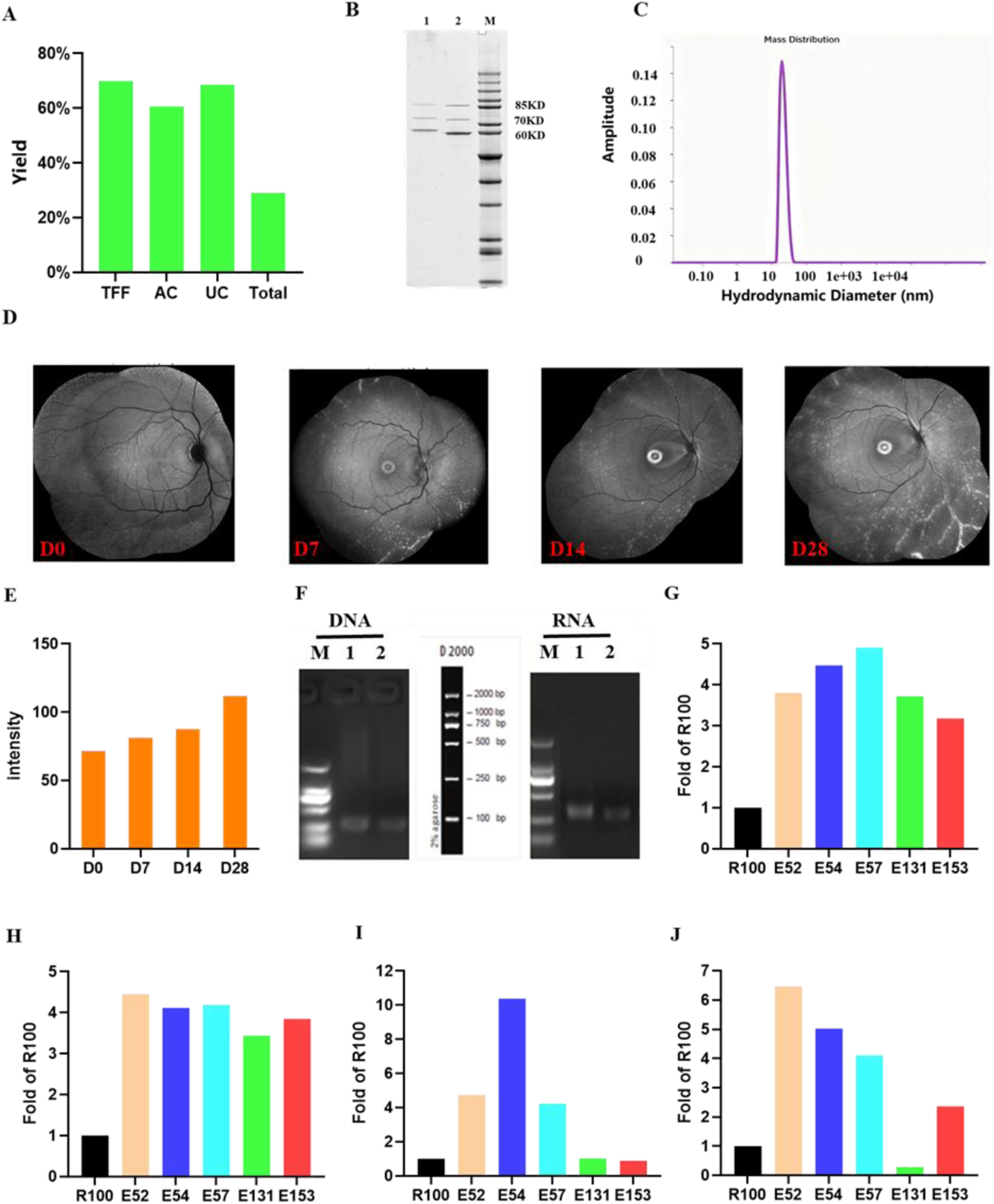
NHP sample preparation and in vivo IVT injection enhanced target cell transduction efficiency. (A) Purification yield of total/each step (relative to the initial sample). Abbreviations: TFF, Tangential Flow Filtration; AC, Affinity Chromatography; UC, Ultracentrifugation. (B) SDS-PAGE image of purified AAV samples. The final samples and AAV reference standard were loaded at the same genome copy number (1E10 vg). Lane 1: purified NHP sample, lane 2: AAV reference standard, lane M indicates protein marker. (C) DLS result of the purified AAV sample. The homogeneity of this final purified NHP sample was analyzed by DLS. (D) BAF imaging was performed on Days 0, 7, 14 and 28 during the NHP study. (E) Fundus GFP Expression Intensity Images in NHP. (F) Pre-experimental DNA and RNA amplification electrophoresis of NHP ocular tissues (M: marker; Lane 1: Neural retina; Lane 2: RPE). (G) AAV distribution data in the NHP RPE. The RNA levels of R100 and engineered variants in the RPE were detected with NGS sequencing. (H) AAV distribution data in the NHP RPE. The DNA levels of R100 and engineered variants in the RPE were detected with NGS sequencing. (I) AAV distribution data in the NHP neural retina. The RNA levels of R100 and engineered variants in the neural retina were detected with NGS sequencing. (J) AAV distribution data in the NHP neural retina. The DNA levels of R100 and engineered variants in the neural retina were detected with NGS sequencing.

Following IVT administration in cynomolgus macaques, blue-light fundus autofluorescence (BAF) imaging from Day 7 showed punctate fluorescence distribution, intensifying by Day 28 (Figure 3D, 3E). We performed preliminary experiment to confirm feasibility of these collected samples, and then these samples were analyzed with NGS sequencing (Figure 3F). NGS analysis of retinal tissues revealed that AAV variant E52, E54, and E57 significantly enhanced transduction efficiency in both RPE and neural retina compared to R100. In the RPE, both the genome copy numbers and RNA level increased 3-5 fold over R100, while in the neural retina samples, these increases reached 5-10 fold (Figure 3G-3J). Of note, variants E131 and E153 demonstrated specific targeting to the RPE, with RNA level 3-4 fold of R100 in RPE while exhibiting only about 50% of R100 levels in the neural retina (Figure 3G-3J). This confirms the enhanced target tissue transduction efficiency of these variants.

### 4. Engineered AAV Variants Show a Favorable Safety Profile in Non-Human Primates

To assess systemic exposure safety, AAV vector biodistribution in major non-target organs was measured via NGS. Following IVT injection, engineered variants showed minimal to undetectable accumulation in the liver, spleen, heart, muscle, kidney, lung, and dorsal root ganglia (DRG). The levels represented only a trace fraction of those observed for intravenously administered AAV9, indicating effective ocular confinement and negligible systemic leakage (Figure 4). The high retinal potency suggests potential for lower clinical doses, further reducing systemic exposure risk.

**Figure 4.**
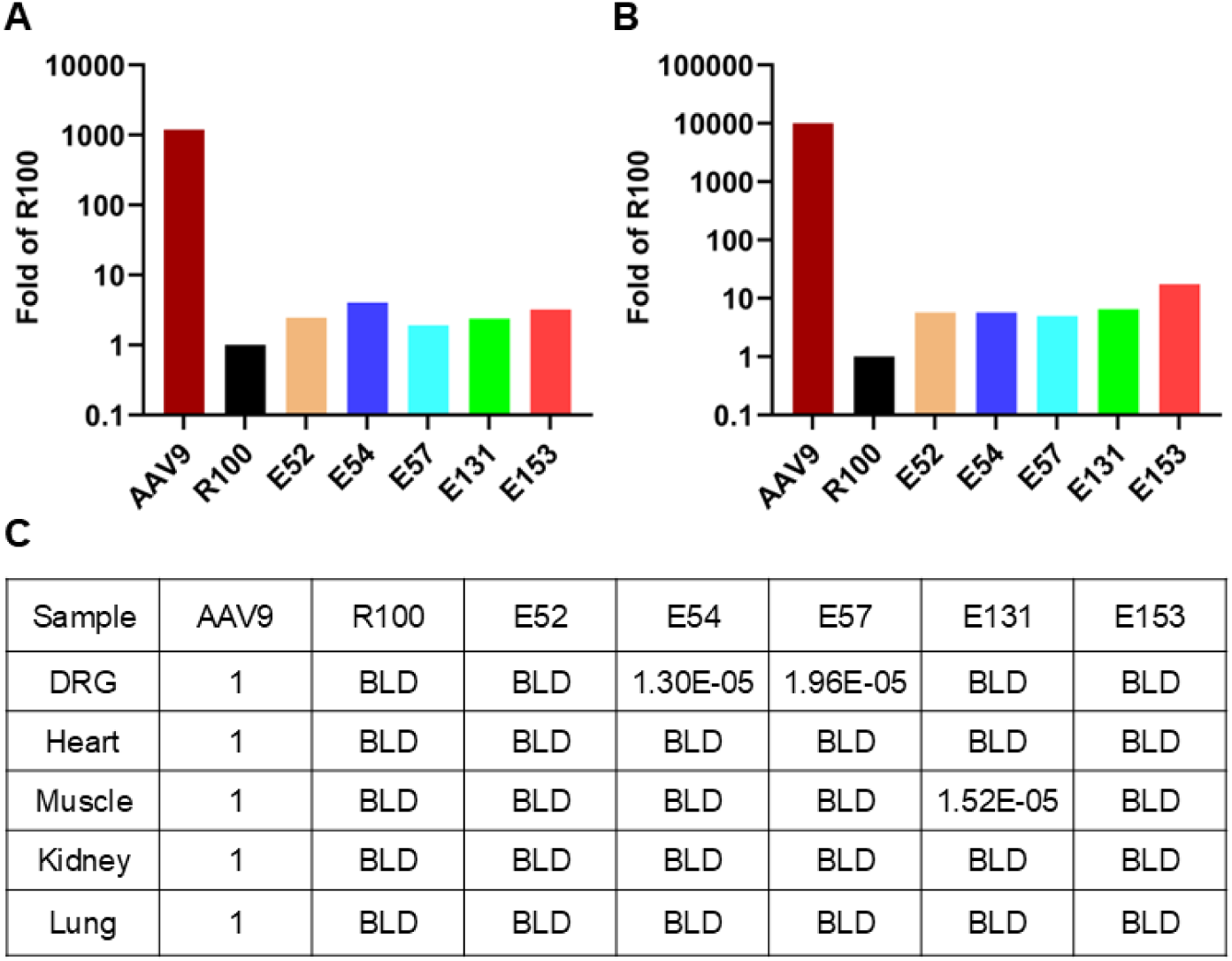
Engineered AAV Variants Show a Favorable Safety Profile in Non-Human Primates. (A) Fold change in AAV distribution in the liver relative to R100. (B) Fold change in AAV distribution in the spleen relative to R100. (C) AAV distribution in DRG, heart, muscle, kidney, and lung relative to AAV9. AAV9 sample was IV injected to NHPs working as a standard, and other AAV variants were IVT injected. Tissues of interest, such as liver, spleen, DRG, heart, muscle, kidney, and lung, were collected and detected for viral genome distribution with NGS sequencing.

### 5. Engineered Variants Demonstrate Excellent Medicine Development Potential

A systematic developability assessment was conducted. Production titers ranged from 1.3×10¹¹to 1×10¹²vg/ml based on our in-house upstream processes (Figure 5A). A multi-step purification process, including TFF, AC and AEX, for variant E54 achieved an overall yield of 28% and a full capsid ratio of 82.2%, which is in the top-tier of this field (Figure 5B-5D). The final product showed high purity (SDS-PAGE), excellent homogeneity (DLS, SEC monomer peak 98.8%), and good stability over multiple freeze-thaw cycles and at room temperature for 72 hours (Figure 5E-5H). In vitro transduction assays confirmed that the variants efficiently mediated protein expression of different genes of interest (GOI), verifying functional activity (Figure 5I).

**Figure 5.**
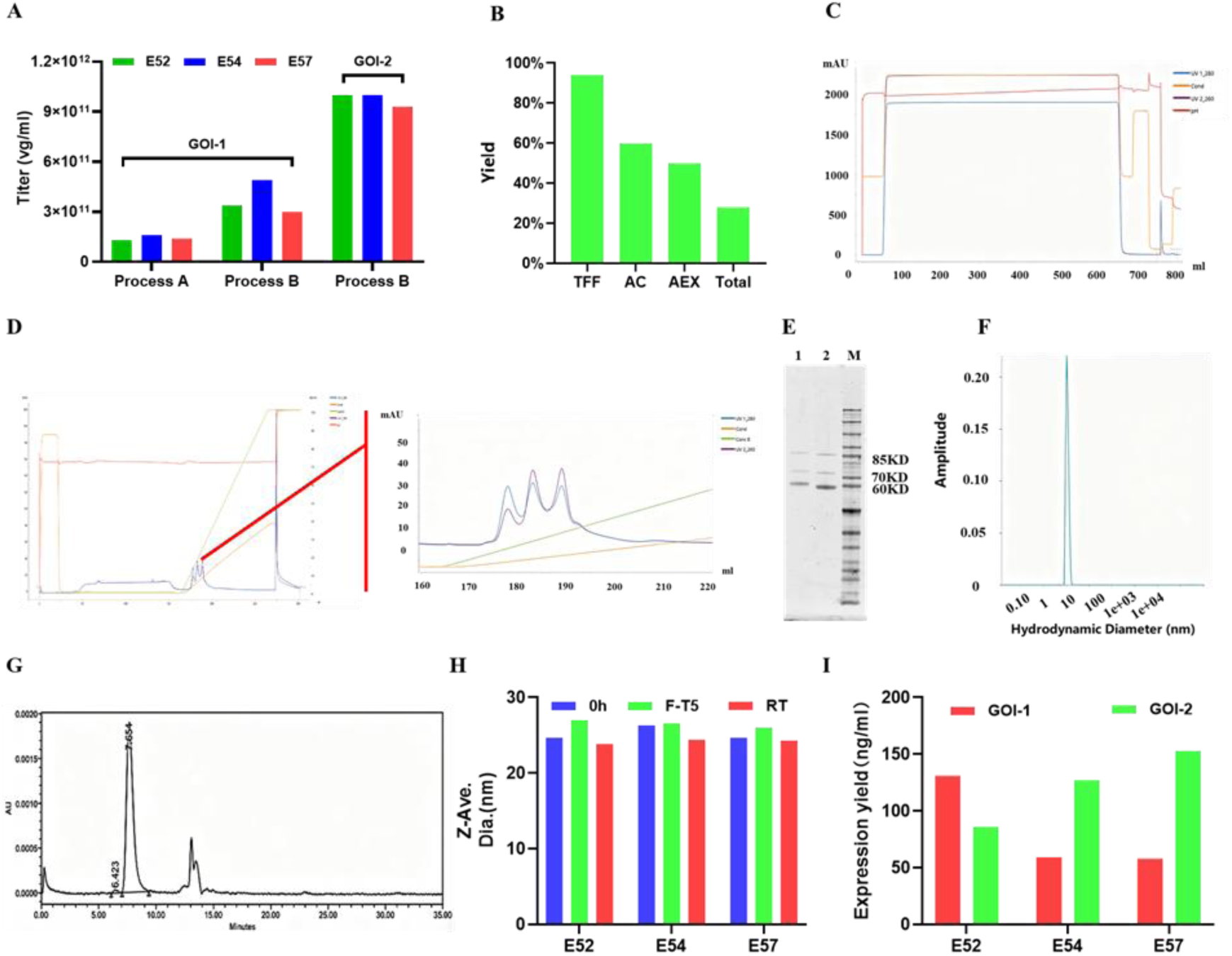
Evaluation of manufacturing process developability for candidate capsid. (A) Virus sample production titer. Virus samples were produced by triple plasmids transfection with HEK293 cells and virus lysate samples were collected 3 days later. The virus titer was detected with qPCR. (B) Purification yield of total/each step (relative to the initial sample). Abbreviations: TFF, Tangential Flow Filtration; AC, Affinity Chromatography; AEX, Anion Exchange Chromatography. (C) Affinity chromatography chromatogram. The E54 samples were concentrated and buffer-exchanged with TFF, and then treated samples were purified with affinity chromatography with AKTA system. (D) AEX chromatogram. The eluted E54 samples from affinity chromatography were further polished with AEX. (E) SDS-PAGE image of purified E54 samples. The final sample and AAV reference standard were loaded at the same genome copy number (1E10 vg). Lane 1: purified E54 sample, lane 2: AAV reference standard, lane M indicates protein marker. (F) DLS result of the purified E54 sample. The homogeneity of this final purified NHP sample was analyzed by DLS. (G) Size-exclusion chromatography (SEC) profile of the purified E54 sample. (H) Particle size distribution of AAV vector under freeze-thaw and room temperature conditions. (I) GOI protein expression driven by AAV transduction in vitro.

## DISCUSSION

This study successfully identified novel AAV variants capable of high-efficiency, specific retinal transduction via IVT injection by combining a rationally designed AAV capsid library, in vitro screening on human RPE cells, and in vivo validation in NHPs. Our key finding is that these candidates mediate transduction levels in the NHP neural retina and RPE significantly surpass current IVT injection benchmark R100 (5-10 fold higher). Importantly, these high-efficiency variants showed nearly undetectable distribution in systemic organs, highlighting exceptional local targeting and potential safety advantages.

Our work directly addresses the clinical challenge of inefficient inner limiting membrane penetration by wild-type AAVs via the IVT route. The initial in vitro screening on ARPE-19 cells enriched for variants with potential human RPE tropism. The subsequent in vivo NHP screening was crucial for validating clinical translational potential due to the anatomical and immunological similarities to humans. The superior performance of our variants over R100, which was itself evolved in NHPs, validates the effectiveness of our REACH platform.

The exceptional safety profile—minimal systemic leakage—is critical for clinical application, as it mitigates risks of immune responses or off-target toxicity.^11^ The combination of high retinal transduction and stringent ocular confinement suggests a high therapeutic index, enabling potentially safer dosing or stronger efficacy.^3,5^

From a translational perspective, these variants hold clear therapeutic potential. The ability to achieve efficient retinal transduction via IVT could enable lower doses or more durable efficacy.^12^ Their favorable manufacturability and stability profiles support commercial scalability.^13–15^ These capsids can serve as universal scaffold vectors for delivering gene therapies for various retinal diseases.

Study limitations include evaluation primarily in healthy NHPs; future work should assess therapeutic efficacy in genetic retinal disease NHP models. Long-term transgene persistence, immunogenicity, and potential retinal pathology require longer-term observation.^16–18^ Although distribution was minimal in major organs, more sensitive assessment of systemic exposure to tissues remains necessary for comprehensive preclinical safety evaluation.^19,20^

In conclusion, we report a class of novel AAV capsid variants that achieve one of the most efficient IVT-mediated retinal transductions reported in NHPs while maintaining high intraocular localization. This work provides promising candidate delivery vectors for retinal gene therapy and validates a platform strategy integrating rational design, human cell model screening, and primate validation for accelerating the development of clinically applicable targeted AAV vectors.

## MATERIALS AND METHODS

### AAV Vector Production and Purification

All AAV vectors (including AAV2, R100, AAV2.7M8, and engineered variants) were produced using the standard triple-plasmid transfection method in suspension HEK293 cells (Thermo Fisher Scientific, VPCs2.0, A52021).

- **Cell Culture and Transfection:** HEK293 cells were maintained in CD05 medium (OPM Biosciences, P688293) and passaged when density reached ≥3×10⁶ cells/mL. For transfection, cells were seeded at 2×10⁶ cells/mL. A mixture of three plasmids—the adenoviral helper plasmid, the AAV rep/cap plasmid (wild-type or variant), and the transgene plasmid encoding enhanced green fluorescent protein (EGFP)—was combined at a 1:1:1 molar ratio. The DNA was complexed with polyethylenimine (PEI) transfection reagent (Yeasen, 40816ES03) at a PEI:DNA=2:1 ratio (v/w) and added to the cells.
- **Harvest and Lysis:** Three days post-transfection, cells were lysed by adding a lysis buffer containing Tris, MgCl₂, and Tween-20, along with Benzonase (50 U/mL, Novoprotein, GMP-1707) to digest unpackaged nucleic acids. The lysate was incubated at 37°C for 3 hours and then clarified by centrifugation.
- **Purification:** The clarified harvest was filtered and subjected to affinity chromatography using an AAVX resin. The eluted virus was further purified by iodixanol (Shanghai Yuanpei, R714JV) density gradient ultracentrifugation. The final viral band was collected, buffer-exchanged, and concentrated using tangential flow filtration (TFF). Viral genome titers were determined by quantitative PCR (qPCR), and the percentage of full capsids was confirmed to be about 70%.
- **AAV Virus Release Test:** The titer of AAV sample was detected qPCR. The particle size distribution was analyzed by dynamic light scattering (DLS). The purity and full-particle ratio were analyzed by sodium dodecyl sulfate-polyacrylamide gel electrophoresis (SDS-PAGE). For the full capsid ratio, an AAV sample (full capsid ratio: about 70%) was used as reference standard. And the percentage of full capsid was calculated based on the protein band intensity ratio measured by Image J.

### In Vitro Transduction Assays

The engineered capsids were screened in vitro using human ARPE19 cells and HEK293 cells.

- **ARPE19 Cell Transduction:** Adult Retinal Pigment Epithelial cell line-19 (ARPE19, ATCC) were seeded in 24-well plates at 1×10⁵ cells/well in DMEM/F12 complete medium (Gibco, 11330032). The next day, cells at 60-80% confluence were infected with AAV vectors at various multiplicities of infection (MOI) in serum-free medium for 4 hours, followed by addition of complete medium to reach 10% FBS. After 48 hours, transduction efficiency was assessed by fluorescence microscopy for GFP expression and quantified by flow cytometer (BD Biosciences) to determine the percentage of GFP-positive cells and mean fluorescence intensity.
- **HEK293 Cell Transduction:** HEK293 cells (Thermo Fisher Scientific, VPCs2.0, A52021) were seeded in 24-well plates at 1×10^6^ cells/ml in CD05 medium (OPM Biosciences, P688293) . Then, the cells were infected by adding viral vectors at different multiplicities of infection (MOI). GFP expression was analyzed 48 hours post-infection via fluorescence microscopy and flow cytometry.

### Large-Scale Production and Evaluation in Non-Human Primates (NHPs)

- **Library Production for NHP Study:** For the NHP study, a pooled library of barcoded AAV variants was produced. The pooled library was purified using TFF, AAVX affinity chromatography, iodixanol density gradient centrifugation, and buffer exchange. The final product was assessed for titer, purity (by SDS-PAGE), capsid full/empty ratio, homogeneity (by DLS), endotoxin level, and sterility to meet administration criteria.
- **NHP Experimental Design:** The study was conducted in cynomolgus macaques (Macaca fascicularis). Prior to dosing, animals were screened for pre-existing neutralizing antibodies (NAbs), and qualified macaques with the lower NAb titers were selected.
- **Dosing and Tissue Processing:** Each animal received a single IVT injection of the pooled AAV library at a dose of 2×10^11^ vg/eye. BAF imaging was performed on monkey eyes using a laser scanning ophthalmoscope on days 0, 7, 14, and 28. 28 days post-dosing, animals were perfused and necropsied. Tissues, including neural retina, RPE, liver, heart, muscle, kidney, and dorsal root ganglia (DRG), were collected and stored at -80°C.
- **Biodistribution Analysis via NGS:** Genomic DNA was extracted from all tissues. The unique barcode was amplified by PCR from each sample, and then subjected to NGS sequencing. The relative abundance of each barcode (corresponding to each AAV variant) was calculated to determine the biodistribution profile.

### Statistical analysis

Unpaired two-tailed t tests were performed in GraphPad Prism and are reported in figures or figure legends. A p value <0.05 was considered significant. * indicates p<0.05, ** indicates p<0.01, and *** indicates P<0.001.

## DATA AVAILABILITY

The data that support the findings of this study are available from the corresponding author upon reasonable request.

## ACKNOWLEDGMENTS

This study was funded by Qilu Pharmaceutical Co., LTD. The authors thank the Institute of Biopharmaceuticals team, pre-clinical team, et al. for their contribution in designing/performing the experiments and data analysis.

## AUTHOR CONTRIBUTIONS

Conceptualization: ZW, YL, ZA; Methodology: ZW, YL, LZ, QZ, PZ, LW, TX; Experimentation: ZW, XX, ZS, HL, QZ, PZ, RH, MY, SW, CH; Resources: ZA, YL, DL, BS; Formal Analysis: ZW, HL, ZS, XX; Investigation: ZW; Writing-Original Draft: ZW, HL, ZS, XX; Writing-Review & Editing: YL, ZA, LZ.

